# Optimizing large organ scale micro computed tomography imaging in pig and human hearts using a novel air-drying technique

**DOI:** 10.1101/2021.07.29.454121

**Authors:** N. Pallares-Lupon, G. Ramlugun, V. Ozenne, J. Duchâteau, A. Delgove, J. Bayer, A. Moreno, M. Constantin, D. Gerneke, G.B. Sands, M.L. Trew, M. Hocini, M. Haissaguerre, E.J. Vigmond, B. Quesson, O. Bernus, R.D. Walton

## Abstract

Underlying electrical propagation in the heart and potentially fatal arrhythmia is the cardiac microstructure. Despite the critical role of muscle architecture, a non-destructive approach to examine not only myocyte orientation, but cellular arrangement in to laminar organization is lacking in hearts from translational animal models and humans. X-ray micro computed tomography using contrast enhancing agents achieves three-dimensional images at near-histological resolutions. However, imaging large mammalian hearts presents challenges including X-ray over-attenuation and loss of image contrast. The goal of this study was to rethink tissue pre-treatment to optimize, and benefit from micro computed tomography imaging resolution in large tissues. Whole pig and human hearts were dehydrated and perfused with a tissue reinforcing agent, hexamethyldisilazane, and slowly air-dried. Heart morphology was conserved and temporally stable. This enabled direct air-mounting for micro computed tomography imaging. Moreover, the desiccated tissue density was significantly reduced compared to the initial hydrated state (P=0.04). Three-dimensional image reconstructions of air-dried hearts segmented using a single intensity threshold revealed detailed microstructural architecture of myolaminae. Conversely, one-step segmentation of hearts loaded with contrast agents poorly estimated the gross anatomical morphology of the heart and lacked identification of tissue microarchitecture. Air-drying large mammalian hearts optimizes X-ray imaging of cardiac microstructure.

## Introduction

Cellular organization in cardiac muscle is a major determinant of electrical propagation in healthy individuals and across the spectrum of electrical disorders. Disruption to the cellular arrangement of cardiac muscle and a biased development of the extracelullar compartment of the heart associates with greater risk of sudden cardiac death(ref). At the microstructural level, reduced cell-to-cell coupling and isolated regions of cardiac myocytes enabling disparate pathways of electrical conduction all promote life-threatening arrhythmia (refs). The scale of structural impairment linked to fatal electrical disorders can be highly localized and below the clinical resolution. For example, Haissaguerre et al. studied young adult patients surviving ventricular fibrillation and absent of clinical signs of structural disease. Despite that, discrete electrical abnormalities were observed at the sites corresponding to the onset of ventricular arrhythmic behavior, which is a hallmark signature of disrupted cellular arrangement (ref - Haissaguerre 2018). Moreover, in patients suffering ischemic insult and chronic structural remodeling, uncoupling of viable but impaired cardiomyocytes from the surrounding myocardium is a pre-requisite for reentrant tachyarrhythmias (refs). Purkinje fibers, forming the ventricular conduction network have also shown to be implicated in reentrant activity (ref – Enjoji 2009; Sinha 2009; Haissaguerre 2002 Lancet). However, precisely distinguishing culprit cell structures and characterizing underlying microstructural abnormalities remains beyond the reach of clinical imaging.

Over recent decades, there has been increased access to large mammalian models of cardiac diseases and human heart tissue. Such specimens yield great potential for scrutinizing the cardiac make-up with higher precision and resolution in the pre-clinical setting. Explanted tissue enables detailed histology, but remains a highly destructive technique and largely two-dimensional. Most studies of electrophysiological behavior based on real tissue geometries and composition rely on magnetic resonance imaging and diffusion tensor methods of local cellular alignment. However, due to partial volume effects, surface cell orientation is often omitted and endocavity structures are often ignored due to challenges in resolving their complex laminar architecture.

X-ray-based micro computed tomography (microCT) is increasingly being applied to image cardiac microstructure^1–6^. Its non-destructive nature, and volumetric information provided, make it a valuable tool. Moreover, the high spatial resolution afforded by microCT, which approaches that of histology (10-25 μm for large field-of-view format systems), provides access to myocyte organization, and even modest, pathologically induced changes in tissue composition^2–4,6^. Yet there is an unmet need for large scale, whole organ imaging of large mammals, including humans. Imaging large tissues tend to come hand-in-hand with compromised working spatial resolutions, dimensionality, and obtainable anatomical detail.

MicroCT of soft tissue, such as the heart, typically requires the use of contrast agents to enhance X-ray attenuation of stained tissue^7,8^. But, increases in sample size come with a concomitant prolongation of the attenuating path length along which X-ray photons need to traverse. The heart is primarily composed of dense laminae composed of a complex arrangement of cardiac myocytes. The major compartments of the heart, the ventricles, are formed out to muscular walls up to 2 cm in thickness along the transmural plane, in humans. But due to the continuous rotation of the X-ray imaging plane, the axial tissue thickness contributing to photon attenuation can be considerably greater. Supplementing tissue with contrast agents, even for low-molecular weight molecules, excessively impacts the coefficient of attenuation of X-rays^9^. Moreover, conventional microCT systems typically produce insufficient X-ray energy for photon detection at this scale of ventricular myocardial samples, which impacts 3D reconstruction and resulting image quality.

MicroCT imaging of explanted fixed samples following treatment with contrast agents is furthermore subject to several constraints. Contrast agent loading is often confounded by diffusion limits, potentially leading to inhomogeneous distribution of the X-ray attenuating molecules^10^. Furthermore, to conserve sample integrity and stability, samples are imaged immersed in the contrast agent solvent that promotes leaching of the contrast agent from the tissue. As sample size increases and X-ray attenuation is greater, the required exposure interval for X-ray photon detection inevitably increases, prolonging the overall acquisition time. Hence, temporal effects of contrast agent mobility become an important factor for larger samples. Moreover, leakage of contrast agents into the surrounding medium reduces the net X-ray attenuation of the sample, compared to the immersion fluid, thereby reducing overall image contrast.

The goal of the study was to achieve micrometric (of the order of 20 μm) images of cardiac structure in intact whole hearts from pigs and human by rendering tissue morphologically stable for microCT imaging directly in air to optimize x-ray attenuation and image contrast. We developed a novel dehydration procedure that eliminated all liquid content of the sample while preserving the overall sample morphology. By doing so, we sought to eliminate the need for X-ray contrast agents, optimize the sample:background contrast, and reduce overall tissue density in large mammalian hearts.

## Materials and Methods

### Tissue samples

Hearts were obtained from pigs (n=6) weighing 20 to 30 kg in accordance with the guidelines from the Directive 2010/63/EU of the European Parliament on the protection of animals used for scientific purposes and the local ethical committee. Procurement and use of a human donor heart with informed consent from family members was approved by the National Biomedical Agency and in a manner conforming to the declaration of Helsinki. The donor hearts was procured at the Bordeaux University Hospital and transported in ice cold cardioplegia to the laboratory. Pigs were premedicated with ketamine (20mg/kg) and acepromazine (0.1 mg/kg), and anaesthesia was induced by propofol (1mg/kg) and maintained under isoflurane, 2%, in air (O_2_) (50%/50%) after intratracheal intubation. Pigs were heparinized after a median sternotomy and before the isolation of the pericardium. Pigs were euthanized by intravenous injection with pentobarbital (15 mL/25 kg). Pig and human explanted hearts were cannulated by the aortic root, and flushed with cardioplegic solution, containing (mmol/L): NaCl, 110; CaCl_2_, 1.2; KCl, 16; MgCl_2_, 16; NaHCO_3_, 10; glucose, 9.01 and heparin, 5000 U/l at 4°C. Macroscopic examination was performed to verify the absence of overt cardiac disease.

### Tissue preparation for microCT

Aortic root cannulas were removed and replaced individually cannulating both left and right coronary ostia for optimal tissue perfusion via the heart’s vasculature. All hearts were pre-treated by re-circulating a perfusate containing phosphate buffered saline (PBS, pH7.4 at room temperature) [Sigma] supplemented with ethylenediaminetetraacetic acid (EDTA) [Sigma], 10 mM and the vasodilator Papaverine (Sigma), 30 μM. Hearts were then perfusion-fixed with formaldehyde, 4% for 2 hours before rinsing three times for 20 minutes in PBS under perfusion. Hearts designated for conventional contrast agent loading (N=2) were perfused with Lugol’s solution (iodine, 5 g/L and potassium iodide, 10 g/L in purified water) for 72 hours. To remove excess contrast agent and fixative, hearts were subsequently washed by agitation in distilled water for 24 hours prior to microCT imaging. Hearts designated for air-drying (pig, N=4 and human, N=1) first underwent serial dehydration using ethanol at increasing concentrations (30, 50, 60, 70, 75, 80, 85, 90, 95, 100, 100%) for a minimum of 1 hour at each step. To prevent tissue deformation during air-drying, coronary vessels were perfused with a 50:50 mix of hexamethyldisilazane (HMDS) [Sigma] and absolute ethanol for 30 minutes at room temperature under a fume hood. This was followed immediately by HMDS, 100% for 2 hours at room temperature. Following HMDS treatment, hearts were hung to air-dry under the fume hood and inside small containers to reduce airflow for 7 days.

### MicroCT imaging and reconstruction

All hearts were imaged using MicroCT (SkyScan 1276, Bruker, Belgium). For contrast-loaded samples, X-ray transmission images were acquired using X-ray source energies of 100kV and 200 μA, which passed through an aluminium and copper plate filter. Imaging of air-dried hearts used optimized parameter settings for X-ray source energies: 55kV and 150 μA and an aluminium plate of thickness 0.5 mm. An average of three acquisitions were captured at rotation steps incrementing by 0.18° over 360°. Total acquisition time was 18 hours per heart. Short axis images were obtained by 3D tomographic reconstruction of the raw images using the cone-beam FDK^11^ algorithm using the NRecon software (Bruker). Images with an isotropic voxel resolution of 21.7 μm were reconstructed using the following parameter settings: Ring artefact reduction, 10 and beam hardening, 25%. Image stacks were exported in 8-bit bitmap format.

### Image processing

Using software Amira, Thermo Fisher Scientific, heart-related structures were manually isolated from non-cardiac objects in reconstructed volumes, such as the sample support, heart container for immersed samples and a positioning clamp fixed to the aorta. Background separation from foreground samples was achieved using an intensity threshold defined by minima separating populations of voxel intensity from volumetric histograms of voxel intensity. The local accumulation of myocytes were estimated based on an assumed cylindrical organisation (Amira, see also ^12^). Textural cylinder correlation analysis was performed on a digitally cropped region traversing the mid left ventricle in the short-axis and covering a depth (longitudinal axis of the heart) of 10 mm from a human heart reconstruction. We assumed cylinder lengths of 1.5 mm, angular sampling of 5° and an outer cylinder radius of 230 μm. A second function, trace correlation lines assessed continuity of highly correlative cylindrical contrast. A minimum distance between center lines of cylinders was 250 μm and a minimum continuity length of 600 μm was applied. Continuity was evaluated in a search cone of length 250 μm, throughout an angle of 37° and using a minimum step size of 10%. Structure tensor analysis was performed as previously described by ^13,14^. Sobel-scharr operators were used to calculate derivatives, which were convolved with a Gaussian kernel (standard deviation of 10 pixels) to obtain the three-dimensional tensor. Eigenvalue decomposition was then performed to find the eigenvector corresponding to the smallest eigenvalue, defining the principal cellular orientation at each voxel. Helix angle was then calculated based on the definition by ^14^. Briefly, an axis was manually defined as the longitudinal axis of the ventricles. The tangential plane was defined as the projection of the longitudinal axis on the voxel. The helix angle was then found by calculating the angle between the projection of the eigenvector onto the tangential plane and the short axis plane.

### Heart morphology and density

Heart morphology and density were compared between the initial hydrated (after heart explantation and initial rinsing) and air-dried states. Maximal heart sample dimensions were recorded consistently along the anterior-posterior, lateral and base-apex axes. Heart samples were also weighed and density was estimated as follows:

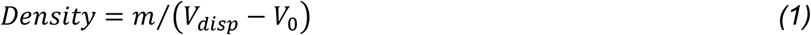

Where *m* is the heart mass (g), *V_0_* is the initial volume (l) of an immersion fluid and *V_disp_* is the displaced volume (l) of an immersion fluid after immersing the heart sample. All statistical comparisons used a Wilcoxon matched-pairs signed rank test with a significance threshold of P=0.05.

### Tissue validation by histology

A short-axis section with a thickness of 5 mm was taken from air-dried ventricles of a pig heart. The tissue section was directly embedded in paraffin over 2 hours at 65°C. Serial sections of 3μm thickness were cut using a microtome (Leica RM2255, France) and mounted on ovesize 50 x 76 mm microscope slides. Sections were treated with toluene to remove paraffin then rehydrated by serial dilutions of ethanol at 100%, 95%, 80%, followed by immersion in tap water. Sections were submitted to Masson’s trichrome staining by immersing sections in xylidine ponceau, 5 mins; rinsing with distilled water, 5 mins; Biebrich Scarlet-acid fuchsin solution, 1 min; 3% phosphomolybdic, 1 min; 1% glacial acetic acid, 1 min; 1% light green SF yellowish, 2 min and then 1% acetic acid, 1 min. Sections were again dehydrated by ethanol 95%, 6 mins, and 100% ethanol, 6 mins, followed by toluene, 3 mins, and stored at room temperature in air. Sections were imaged using an Axio Scan.Z1 slide scanner (Zeiss, Germany).

## Results

### Heart structure

Explanted hearts had a mean wet mass of 173.1±27.1 g. The maximum dimensions of hearts were measured in the longitudinal (base-apex) and short (lateral and anterior-posterior directions) axes, which were 9.2±0.9 cm, 7.7±0.3 cm and 5.7±0.6 cm, respectively. The density of wet heart samples was 1.03±0.03 g/l. However, after air drying heart samples pretreated with HMDS, an overall 2-fold reduction of heart density to 0.54±0.02 g/l was achieved (P=0.04, N=3). This was concurrent with a reduced mass (22.1±6.4 g), yet maximal axial heart dimensions were unaltered (8.9±0.9 cm, 7.0±0.0 cm and 5.8±0.3 cm, respectively). As such, air-dried hearts showed conserved anatomical features (Figure 1). The ventricles retained a morphology consistent with a diastolic phase of contraction. The prominent atrial appendages of pig hearts did not show collapsing of free walls, despite being relatively thin (~1-3 mm).

**Figure1.**
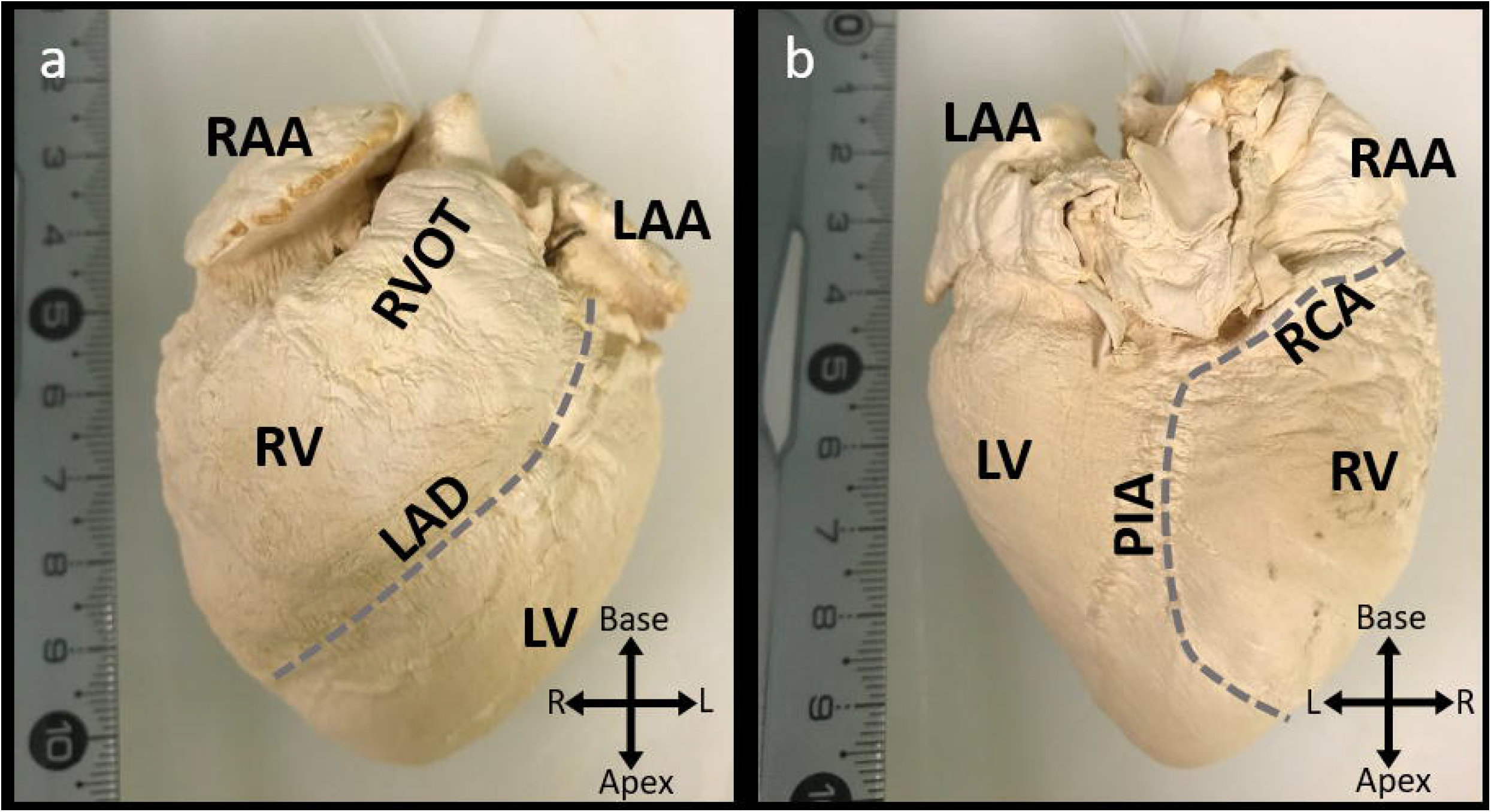
The air-dried pig heart. An example air-dried pig heart is shown in an anterior **(a)** and posterior **(b)** perspective. Abbreviations are: RAA, Right atrial appendage; LAA, left atrial appendage; RVOT, right ventricular outflow tract; RV, right ventricle; LV, left ventricle; LAD, left anterior descending artery; RCA, right coronary artery and PIA, posterior interventricular artery.

### X-ray transmission of the intact heart

X-ray transmission of pig hearts prepared conventionally by staining with iodine contrast agent was impeded by almost complete attenuation (minimum transmission of 6.5%± 1.3%) and a low dynamic range throughout the heart (Figure 2a). By contrast, the air-dried heart maximally reduced transmission to 32.4%± 2.7% (Figure 2b). Moreover, X-ray attenuation throughout the heart sample was relatively heterogeneous for the air-dried heart when compared to the iodine-stained sample. The heterogeneous transmission of X-rays in the air-dried heart related to structural variations and enabled distinction of gross anatomical landmarks such as ventricular cavities, septal regions, atrial appendages, and left and right outflow tracts. Conversely, the ventricles were practically indistinguishable for the iodine-stained heart when examining X-ray projection images. This indicated that X-ray attenuation was, therefore, dependent upon sample structural heterogeneities for the air-dried heart, but sample thickness and the axial X-ray path length were greater determinants of X-ray attenuation for the iodine-stained pig heart.

**Figure 2.**
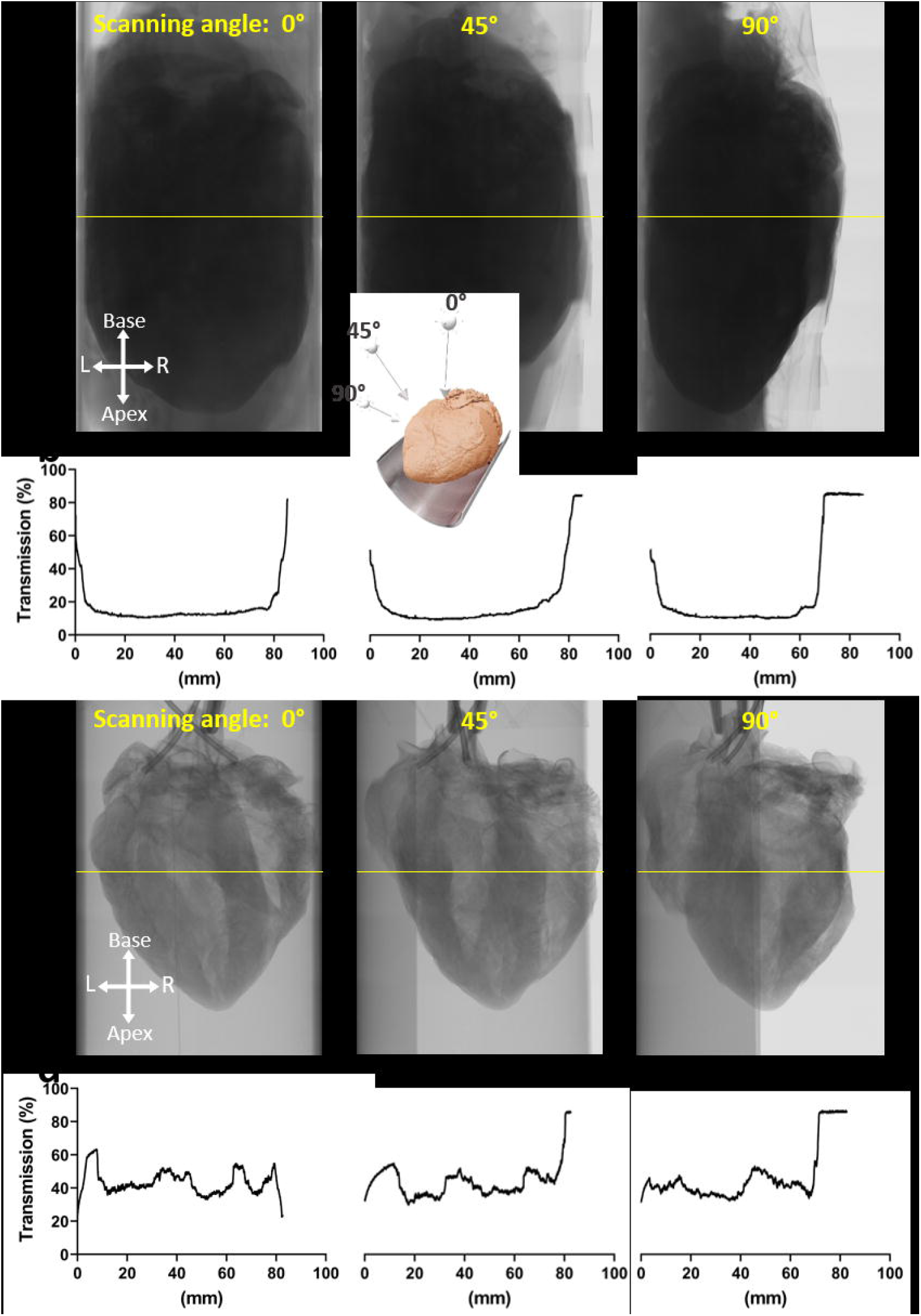
X-ray transmission of the pig heart. **a)** X-ray projections showing transmission through a pig heart prepared with iodine contrast agent and immersed in water. Images are shown for 0°, 45° and 90° of axial rotation, as demonstrated in the inset image. **b)** X-ray transmission profiles were extracted along the heart’s short axis at the yellow lines shown in **a**. **c)** As for A, X-ray projections are shown for an air-dried pig heart. **d)** X-ray transmission profiles extracted along yellow lines in **c**.

### Correlating X-ray attenuation and tissue thickness

X-ray attenuation was correlated with axial tissue thickness of samples dissected from the left ventricular free wall. Figure 3a shows the samples with varying axial thicknesses as three-dimensional volume rendered images used for analysis. Tissue samples were aligned in the axial plane with the corresponding X-ray projection. Here we compared the X-ray attenuation against the axial sample thickness for iodine-stained and air-dried samples. Furthermore, to elucidate the differential impact of the presence of a contrast agent and the immersion fluid, a third sample was air-dried and stained with the iodinated contrast agent (and allowed to air-dry again). Figure 3b shows that sample thickness positively correlates with X-ray attenuation in all conditions. A linear model of the non-dehydrated iodine sample thickness was a poor estimate of X-ray attenuation (R^2^=0.34). An improved linear correlation (R^2^=0.51) was observed when air-drying. Furthermore, it was estimated that 90% of X-ray attenuation would necessitate an air-dried sample thickness of 218.7 mm, vastly improving the dynamic range of compatible sample sizes. Supplementing the air-dried sample with iodine further ameliorated the attenuation-thickness relationship (R^2^=0.58) but yielded a dramatically increased slope. Despite that, the initial attenuation level was reduced by 43.5% compared to the non-dehydrated iodine-stained sample due to the absence of an immersion fluid. As a result, 90% attenuation was estimated for samples of 24.5 mm thickness.

**Figure 3.**
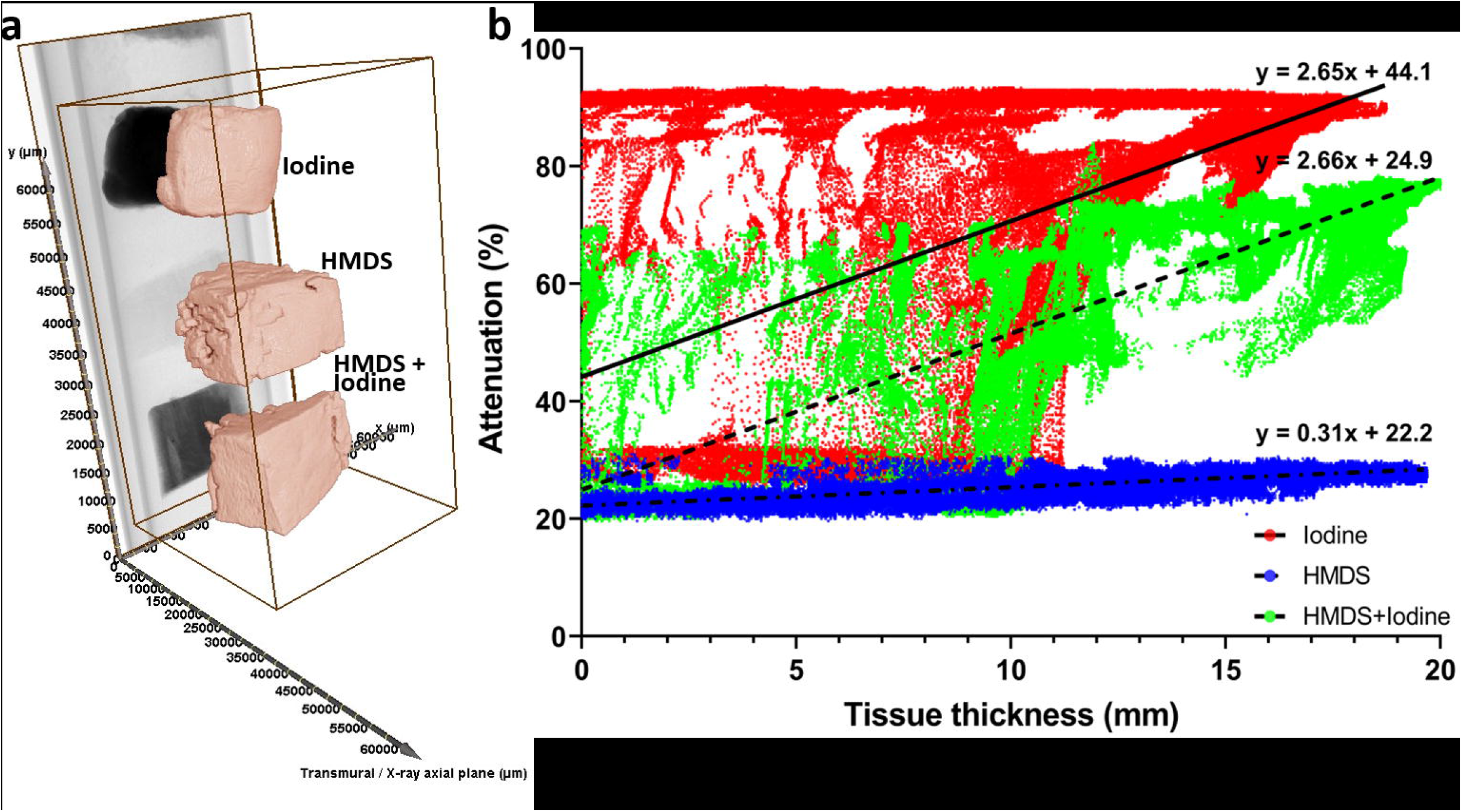
Correlation of sample thickness and X-ray attenuation. Three dissected samples of ventricular tissue were treated with iodine, air-drying and air-drying followed by iodine-staining. **a)** Sample morphology position is rendered in 3D. An X-ray projection image of the three samples is plotted in alignment along the X-ray axial plane. **b)** Sample thickness was linearly correlated with X-ray attenuation, derived from the X-ray projection image plotted in **a**.

### 3D image reconstruction and segmentation

X-ray transmission images of pig hearts were tomographically reconstructed in 3D. Figure 4a shows an example cross-section from an iodine-stained heart. Differential X-ray attenuation was observed throughout the sample as a transmural gradient with the highest intensity signals at the epicardium and decreasing to the endocardium. A low image contrast yield was observed intramurally, and large morphological features such as the epicardial surface and ventricular cavities were distinguishable. However, the immersion medium was also detectable with contrast in the same range as the sub-endocardium. Conversely, the airdried heart sample showed distinct intensity differences between the sample and the surrounding air medium (figure 4b). As a result, a considerably improved morphological representation of the anatomical components of the ventricles was observed. A voxel intensity histogram of the full 3D reconstructed images revealed distinct frequency distributions for iodine-stained and air-dried hearts (figure 4c). Low voxel intensity populations corresponding to air and the background noise levels, and high intensity sample populations, were separated by frequency minima. Yet the intensity distribution minima differed (intensity values of 34 and 45) and the iodine-stained sample showed two peaks in the sample population, but only a single peak was found for the air-dried heart image. The difference in area under the frequency distribution curves is a consequence of non-cardiac (background) voxels contributing to the histogram in the iodine-stained heart image compared to air-drying. Using the respective minima as intensity threshold values, image segmentation was performed. Figure 4d shows the region and morphology of the image surpassing the intensity threshold for iodine. The gross structure of the heart was poorly represented, predominantly due to false detection of heart tissue where immersion fluid was located. Most strikingly, this segmentation approach failed to even distinguish the ventricular cavities. In contrast, the air-dried heart showed excellent correspondence between the segmented region and the heart tissue (figure 4e). Moreover, this rudimentary segmentation approach readily distinguished intramural features including myolaminae, cleavage planes (separations of laminae) and coronary vessels. A 3D representation of the segmented greyscale image of an iodine-stained heart was highly perturbed by the immersion fluid (figure 5a), which gave a low visibility to the far (based on the reader’s perspective) portions of the sample. Moreover, the non-rigid heart was susceptible to assuming the shape of the sample support, as can be observed from parallel lateral borders of the heart in the 3D rendered image. Conversely, the air-dried heart volume was rendered unimpaired in the absence of further image processing (figure 5b). The sample’s rigid structure fully retains the tissue morphology in a diastolic state. From an intracavity perspective, major endocavity structures were clearly identifiable, including the papillary muscles, chordae tendineae, atrio-ventricular valves and trabeculae. Moreover, microstructural components could also be resolved, including free-running Purkinje fibers and intramural muscle laminar structures.

**Figure 4.**
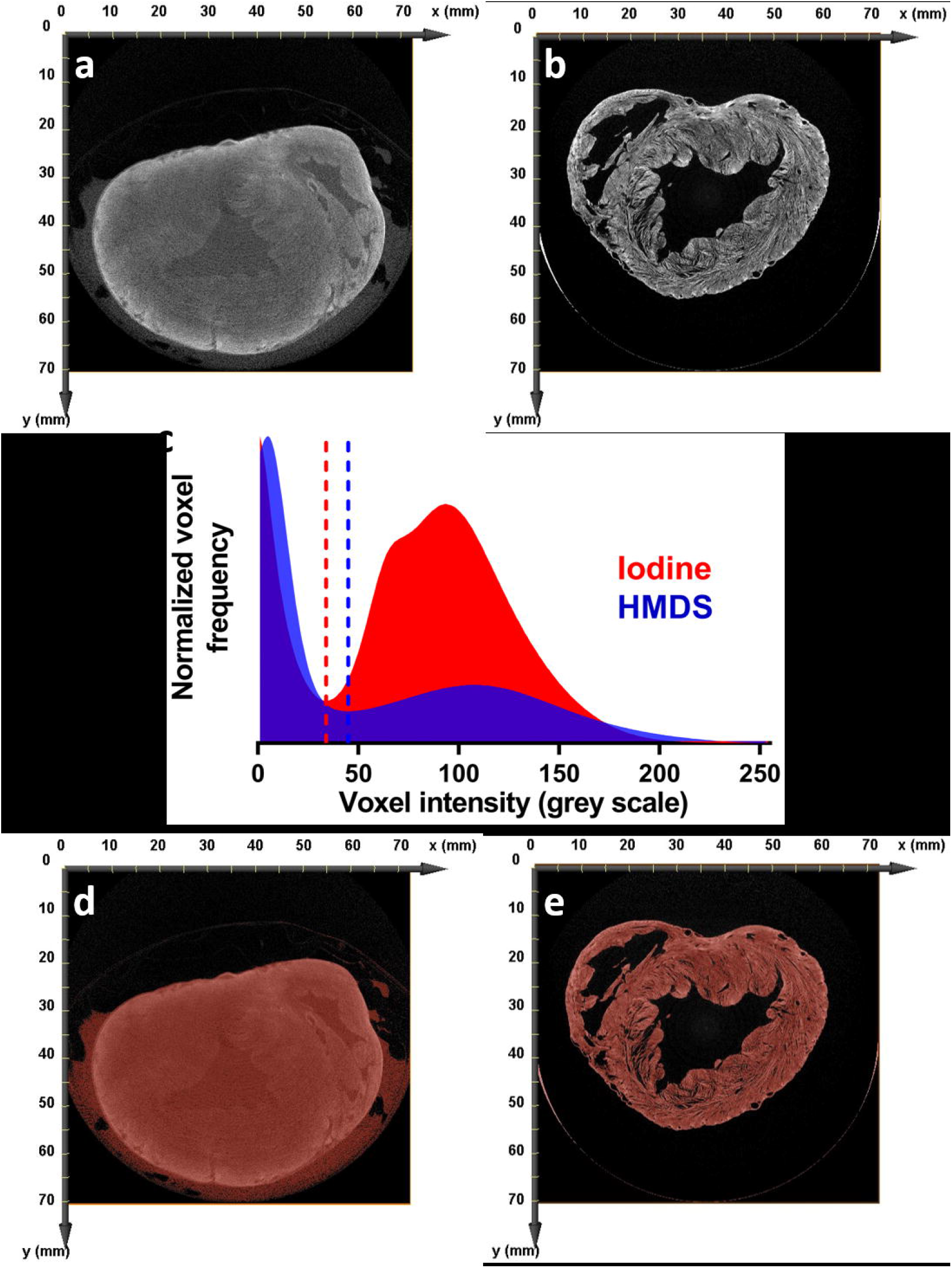
Image volume reconstruction and segmentation of pig hearts. Tomographically reconstructed image slices of iodine-stained **(a)** and air-dried **(b)** hearts in the short-axis. **c)** Voxel intensity distributions for iodine-stained and air-dried (HMDS-treated) hearts. Intensity minima are shown by vertical dashed lines. Arrows indicate voxel intensity population maxima. Intensity thresholds were applied for image segmentation of iodine-stained **(d)** and air-dried hearts **(e)**.

**Figure 5.**
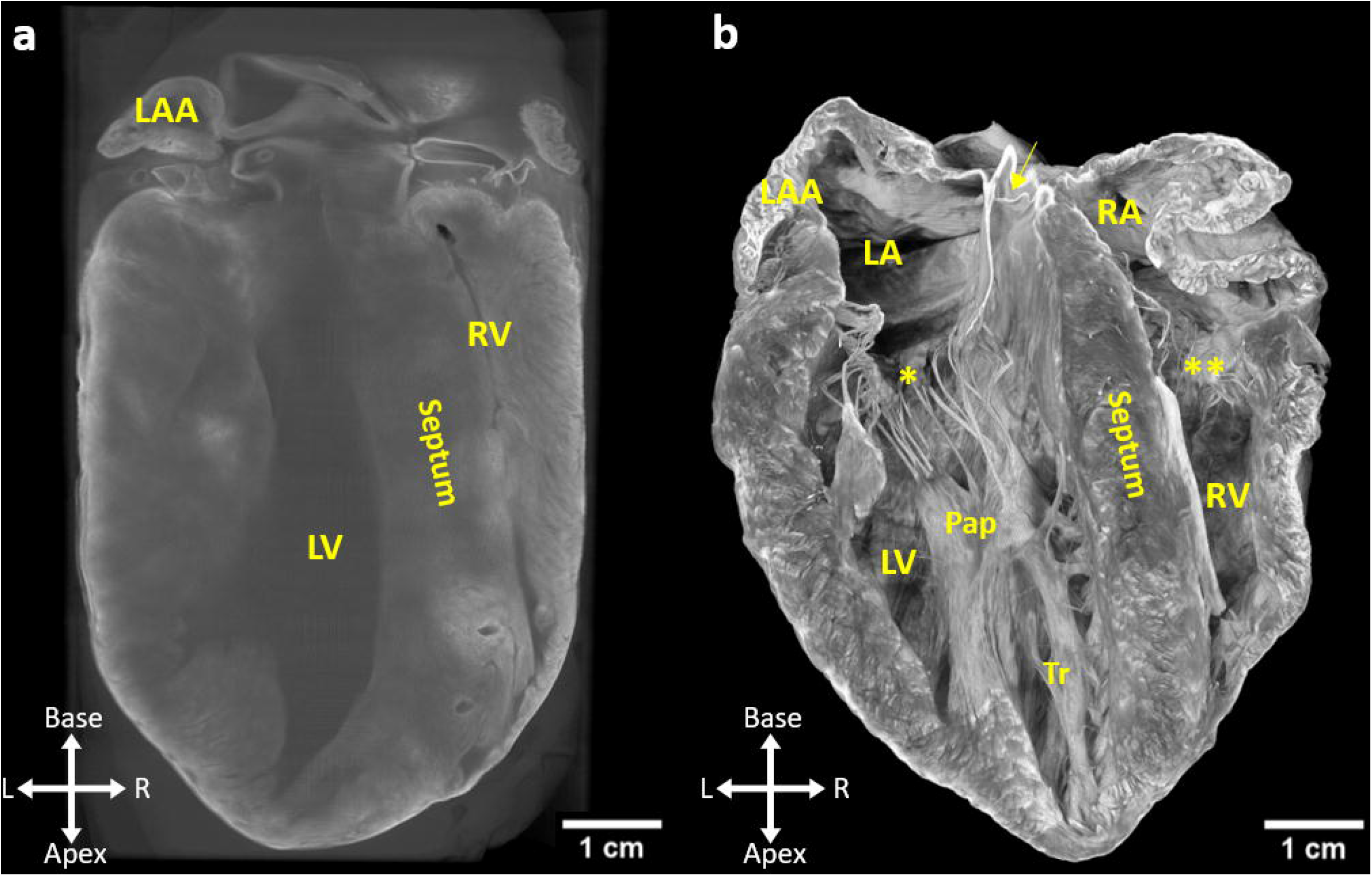
Volume rendered images of iodine-stained (a) and air-dried (b) pig hearts.

### Histological validation

Due to the dehydrated state of the air-dried hearts, samples were compatible with direct immersion in, and infiltration by melted paraffin. In this way, tissue sections for histology could be collected without further deformation of the overall heart sample. Tissue sections could then be labelled with Masson’s trichrome stains. Figure 6a shows that myocardial and collagen-rich compartments and their morphology were retained during the air-drying procedure. For example, ventricular cavities maintained a state of relaxation (diastole) and, at a microstructural scale, the coronary vasculature was not collapsed. Histological sections could be directly compared with microCT images of the same heart sample. Figure 6b shows an orthogonal plane through the reconstructed microCT volume matching the slice orientation and depth of the histological cut. The reconstructed image slice shows consistent myocardial fiber orientation, distribution of connective tissue and vasculature.

**Figure 6.**
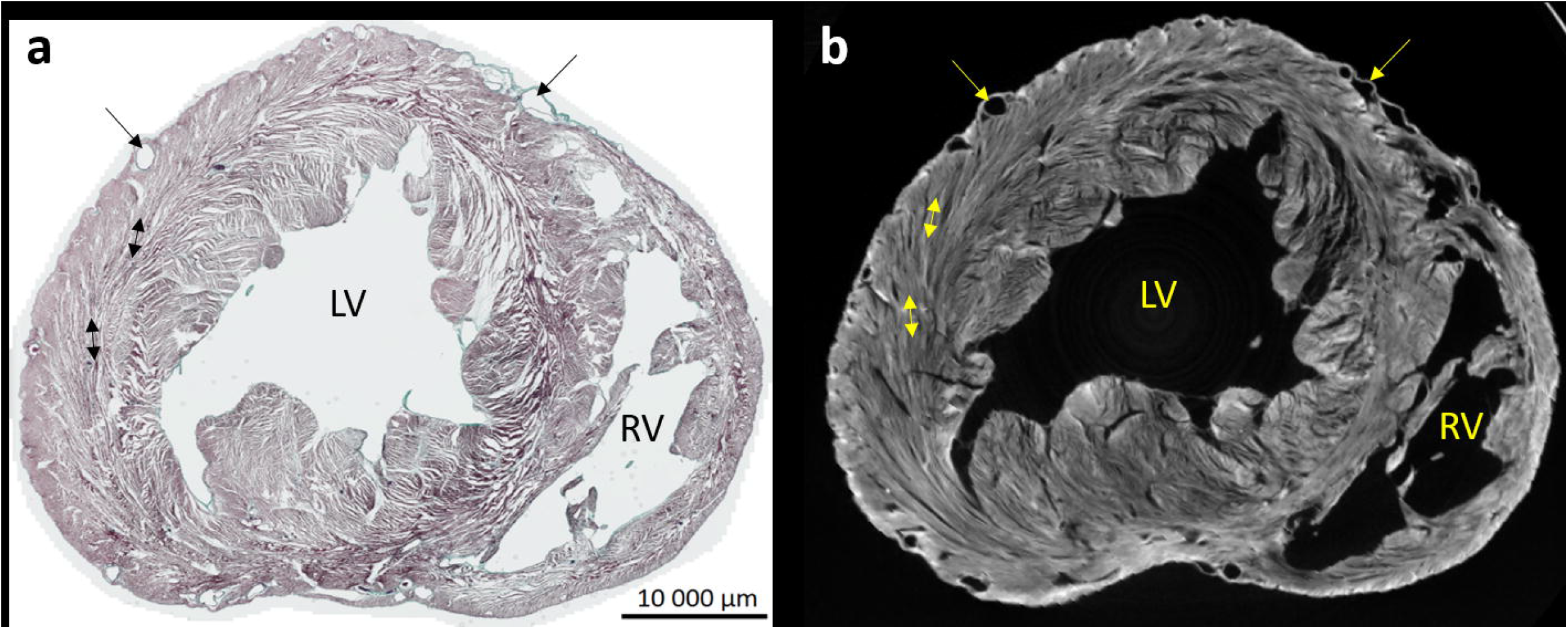
Histological validation of air-dried pig ventricular morphology and composition. **a)** Histological tissue section treated with Masson’s trichrome staining after air-drying. **b)** An image slice was extracted from the volume reconstruction in the same oblique plane as the histological section in **a**.

### Air-dried human heart

The applicability of the dehydration and air-drying approach was evaluated in a human heart. The whole human heart stabilized morphologically in an apparently diastolic state without identifiable abnormal deformation. X-ray projections were comparable to air-dried pig hearts, with minimum X-ray transmissions of 27.2±2.8% measured from 2000 projection images acquired throughout 360° of axial rotation. Tomographic 3D reconstruction of the human heart image presented voxel intensity distributions with two distinct populations (figure 7a). The minimum separating populations served as an effective intensity threshold for segmenting the heart sample (figure 7b). Figure 7c shows a volume rendering of the segmented ventricles. As for the pig (figure 4b), the 3D representation of the human heart reveals easily resolvable endocavity structures, as well as intramural myocyte orientation and laminar organization. We sought to evaluate local cellular orientation in the air-dried human left ventricle. Figure 8a shows a cross-section through the short axis of the left ventricle cut to a thickness of 20 mm in the longitudinal axis. Raw grey scale images show that the crosssection was taken at the level of the insertion of the papillary muscles into the left ventricular free wall (figure 8b), and that the sample was predominantly composed of working myocardium. The helix angle, based on eigenvectors of local image contrast, shows a consistent transmural (epicardium to endocardium) low to high helix angle transition throughout the left ventricle tissue volume (figure 8c). The principal three-dimensional directions of locally aligned myocardium were sought by comparing image intensity profiles to a cylinderical template. Figures 8d & 8e show the cylinder correlation distribution with peaks aligning along the organizing centers of commonly orientated myocytes. Removing voxels containing correlation factors inferior to 45 revealed the myocardial-specific image field (figure 8f). The constrained correlation field was used to trace along the principal orientations of myocytes from seeding points defined by correlation factors superior to 68 (figure 8g). Figure 8h shows the principal cellular alignment throughout the tissue section (figure 8h). Helix angles identified along voxels intersecting traces of cellular alignment showed that the helix angle distribution ranged predominantly from 30° to 150° (figure 8i).

**Figure 7.**
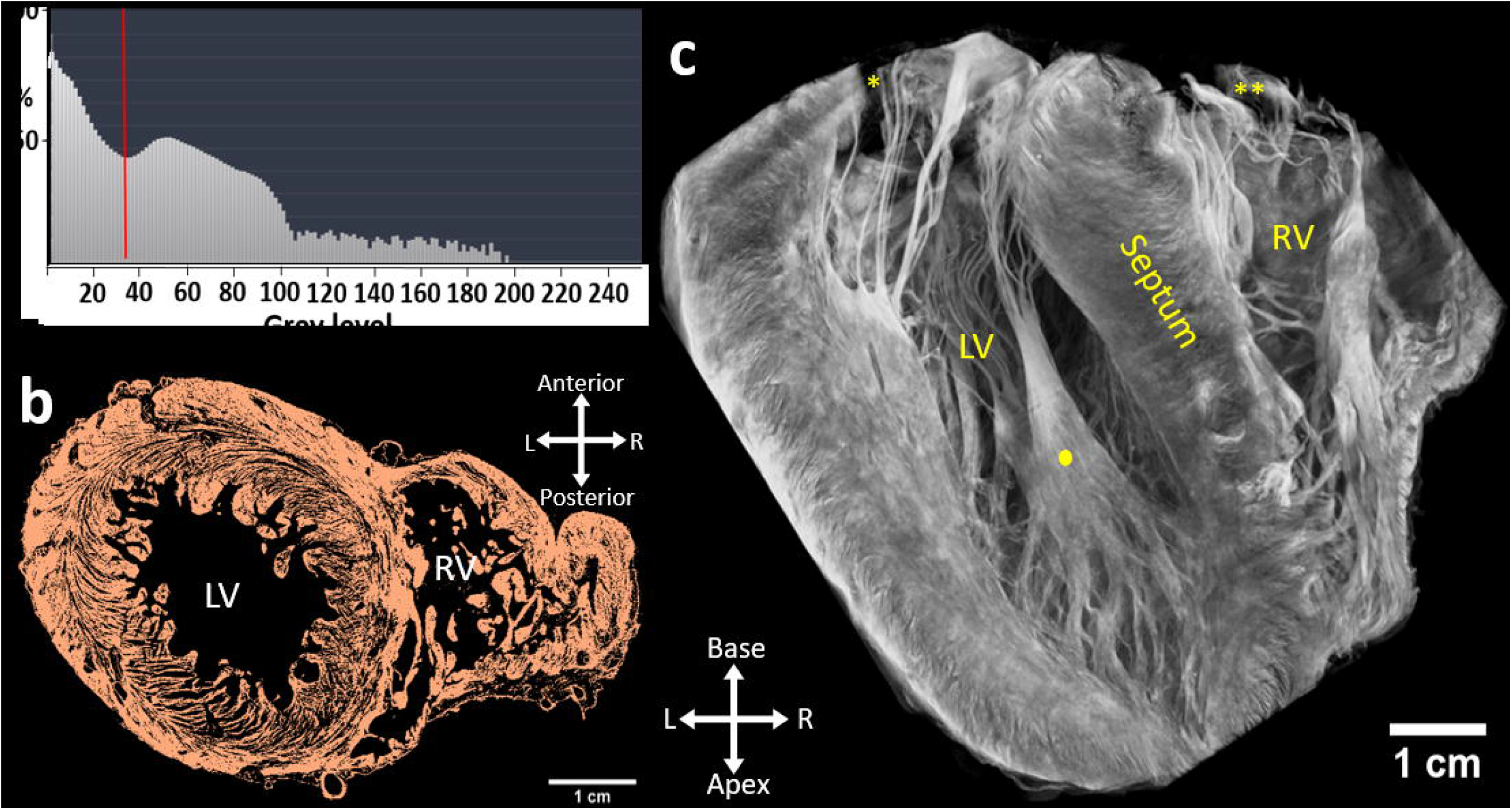
MicroCT image reconstruction of air-dried human ventricles. **a)** A normalized voxel intensity distribution of the image volume. **b)** An orthogonal image slice through both left and right ventricles of the human heart. The region of tissue segmentation by voxel intensity minimum (vertical red line in **a**) is shown in red. **c)** Volume rendering of the segmented ventricles.

**Figure 8.**
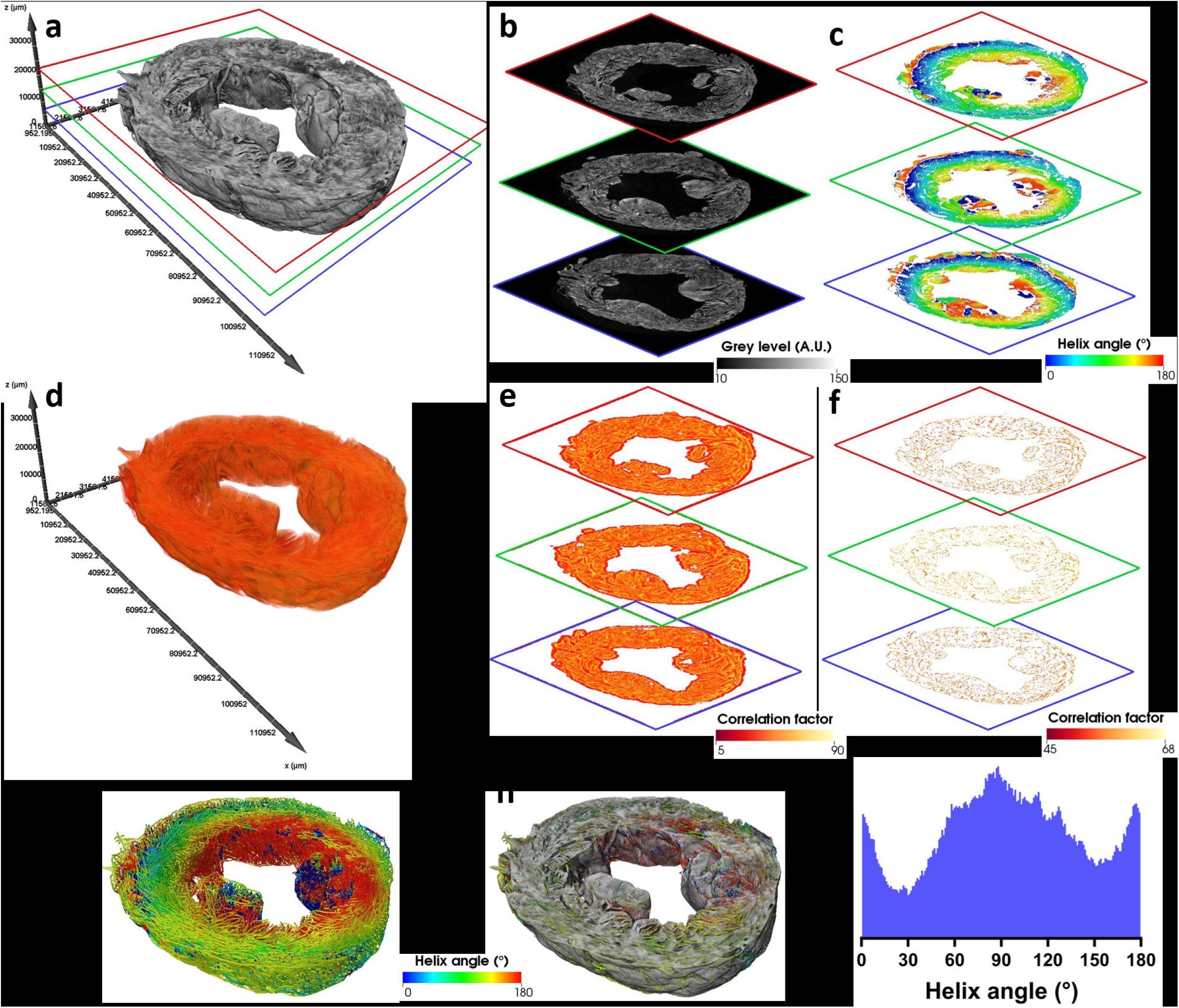
Left ventricular fiber orientation in a human heart. **a)** Volume rendered cross-sectional wedge from a left ventricle. **b)** Image slices from three levels indicated by bordering colors blue, green and red, which correspond to color-matched slice positions in **a**. **c)** Helix angles of each pixel in the segmented tissue region. **d)** Cylinder correlation volume rendering. **e)** Image slices of the cylinder correlation. **f)** Correlation factor images with minimum correlation factor threshold at 45. **g)** Myocardial fiber bundles traced from the reduced cylinder correlation image. Fibers are colored by their local helix angle. **h)** Myocardial fibers superimposed with the greyscale image volume shown in **a**. **i)** Helix angle frequency distribution.

## Discussion

A novel technique was shown to robustly and reproducibly dehydrate and dry pig and human hearts while conserving morphological state. Air-dried tissue was shown to be morphologically stable in air under ambient conditions, enabling long duration microCT scanning without the use of immersion media. A complete removal of moisture from heart samples lowered the overall tissue density, which favorably reduced X-ray attenuation. Moreover, air-drying enhanced the heterogeneity of attenuation within the tissue sample, whereby vascular lumina and interlaminar clefts formed air-gaps. This maximally optimized X-ray attenuation gradients forming between tissue and air, enhancing the image contrast of the detected X-ray transmission within the sample, when compared to conventional contrast agent-based approaches. Tomographic 3D reconstructions of air-dried hearts revealed highly resolvable heart anatomy. A single intensity threshold easily segmented the heart tissue from the background, without further imaging processing. Segmented heart images enabled detailed exploration of the tissue composition and architecture, both at the heart surface and at the microstructural scale below the tissue surface, including laminae and local cellular orientation.

### Air-drying methodology

Air-drying of biological samples has previously been limited to tissue sections in the millimetric and sub-millimetric range, typically for electron microscopic imaging^15^. The major challenge to air-drying samples has been to prevent tissue deformation caused by the surface tension of the evaporating immersion medium acting at the molecular level. The most common method applied for complete dehydration and drying of tissue is critical point drying. This approach relies on expensive, specialized apparatus to replace the sample’s liquid composition with liquid carbon dioxide, which is then evaporated from the sample under a tightly regulated environment to minimize the liquid surface tension^16,17^. Scalability is therefore a limiting factor for critical point drying. An alternative strategy using a chemical reinforcing agent by silylation using hexamethyldisilazane was found to adequately and comparably conserve small sample morphology after just a few minutes of immersion followed by airdrying on the lab bench^18,19^. When considering large tissue samples such as whole organs from large mammals, the diffusion coefficient in tissue impedes uniform exchanges of solutions and temperature gradients. In our study, the coronary vasculature of the heart provided a mechanism for solution access to deeper tissue layers. Therefore, continuous arterial-perfusion, ambient temperatures, and long exposures to perfusates in explanted tissue, facilitated effective serial-dehydration by alcohols and hexamethyldisilazane treatment. During the drying phase, evaporation was slowed by isolating samples from convection airflow. This served to limit the variability of the rate of tissue desiccation, particularly between the exposed epicardial surfaces and deeper muscle layers. Moreover, slowing evaporation further reduced the number of molecules dispersing simultaneously, in turn reducing cumulative forces that evaporating molecules could impose on the tissue’s microstructure. Overall, air-drying whole pig hearts observed an average size reduction of 3.4% (N=3, P=0.25), which is comparable to, or less than shrinkage by formalin fixation or alcohol dehydration observed in other tissues^20,21^.

### Cross-compatibility of air-dried tissues

We observed that air-dried myocardial samples were compatible with subsequent immersion in contrast agents (figure 5). As expected, supplementing tissue with contrast agents increased X-ray attenuation, but, the absence of an immersion medium favorably offset the tissue thickness – attenuation relationship to offer an intermediate level of X-ray attenuation. Moreover, X-ray contrast agents can emphasize heterogeneities of tissue composition based on differential loading^22,23^. Consequently, a combined application of airdrying tissue and contrast agent loading should augment the dynamic range of image contrast. In this study we implemented iodine as the most common contrast agent utilized in soft tissue microCT imaging. Previous studies have shown alternative contrast agents for microCT imaging in hydrated cardiac tissue, such as phosphomolybdic acid and phosphotungstic acid^24^. Further comparisons should be investigated for compatibility and suitability for use with dehydrated and air-dried tissue.

To evaluate the effectiveness of the air-drying method to provide sharp image contrast of large mammalian heart tissue, the local preferential orientation of muscle cells were shown. The myocardium has a complex arrangement of myocytes which is vital for heart function, in terms of the spread of electrical activation, the distribution of contractile force, and the passive compliance of the chambers. Abrupt changes of myolaminar orientation across the ventricular wall are well documented^25^, yet previous studies mapping cellular orientation at the organ scale of large mammals have been limited to image resolutions insufficient to resolve myocardial architecture at the scale of myolaminar thickness (4-6 myocytes in the transverse plane)^13^. We found that microCT of air-dried tissue readily identified individual muscle lamina and enabled extrapolation of their local cellular arrangement using a 3D structure tensor approach^26^. Minimal image processing by applying a single intensity threshold effectively separated laminae. Furthermore, structure tensor analysis of the raw reconstructed volumes enabled procurement of the helix angle distribution throughout the heart sample. Using a cylinder template to identify fibrous textures, it was also shown that the raw image served as an effective input for tracking the continuity of the principal cellular orientation in 3D space.

### Conclusion

In conclusion, preparation of pig or human hearts by dehydration and chemical reinforcement by hexamethyldisilazane effectively stabilizes the tissue for air-drying. Directly mounting tissue in air under microCT optimizes X-ray attenuation of tissue for high contrast and high quality 3D reconstructions. Air-drying of tissue removes the necessity to supplement samples with contrast agents. Moreover, air-dried tissue is highly compatible with performing morphologically-matching histological validation. MicroCT imaging of airdried whole hearts from large mammals enables detailed and quantitative analysis of tissue morphology at the microstructural level in a non-destructive manner.

## Author contributions

RDW conceived and designed the study;

NP-L, GR, JD, AD, JB, AM, MC VO, DG, GBS, MLT & RDW analyzed and interpreted the data;

NP-L, GR, JB, DG, GBS, MLT, MH, EJV, BQ, OB & RDW drafted or critically revised the manuscript;

All authors gave final approval of the submitted manuscript.

## References

1. van Deel, E., Ridwan, Y., van Vliet, J. N., Belenkov, S. & Essers, J. In vivo quantitative assessment of myocardial structure, function, perfusion and viability using cardiac micro-computed tomography. J Vis Exp (2016). doi:10.3791/53603

2. Stephenson, R. S. et al. High-Resolution Contrast-Enhanced Micro-Computed Tomography to Identify the Cardiac Conduction System in Congenitally Malformed Hearts: Valuable Insight From a Hospital Archive. JACC Cardiovasc Imaging (2018). doi:10.1016/j.jcmg.2018.05.016

3. Novo Matos, J. et al. Micro-computed tomography (micro-CT) for the assessment of myocardial disarray, fibrosis and ventricular mass in a feline model of hypertrophic cardiomyopathy. Sci Rep (2020). doi:10.1038/s41598-020-76809-5

4. Detombe, S. A. et al. Longitudinal follow-up of cardiac structure and functional changes in an infarct mouse model using retrospectively gated micro-computed tomography. Invest Radiol (2008). doi:10.1097/RLI.0b013e3181727519

5. Butcher, J. T., Sedmera, D., Guldberg, R. E. & Markwald, R. R. Quantitative volumetric analysis of cardiac morphogenesis assessed through micro-computed tomography. Dev Dyn (2007). doi:10.1002/dvdy.20962

6. Badea, C. T. et al. Cardiac micro-computed tomography for morphological and functional phenotyping of muscle LIM protein null mice. Mol Imaging (2007). doi:10.2310/7290.2007.00022

7. Aslanidi, O. V. et al. Application of micro-computed tomography with iodine staining to cardiac imaging, segmentation, and computational model development. IEEE Trans Med Imaging 32, 8–17 (2013).

8. Stephenson, R. S. et al. Contrast enhanced micro-computed tomography resolves the 3-dimensional morphology of the cardiac conduction system in mammalian hearts. PLoS One 7, (2012).

9. Ashton, J. R., West, J. L. & Badea, C. T. In vivo small animal micro-CT using nanoparticle contrast agents. Frontiers in Pharmacology (2015). doi:10.3389/fphar.2015.00256

10. Li, Z., Clarke, J. A., Ketcham, R. A., Colbert, M. W. & Yan, F. An investigation of the efficacy and mechanism of contrast-enhanced X-ray Computed Tomography utilizing iodine for large specimens through experimental and simulation approaches. BMC Physiol (2015). doi:10.1186/s12899-015-0019-3

11. Feldkamp, L. A., Davis, L. C. & Kress, J. W. Practical cone-beam algorithm. J Opt Soc Am A (1984). doi:10.1364/josaa.1.000612

12. Weber, B. et al. Automated tracing of microtubules in electron tomograms of plastic embedded samples of Caenorhabditis elegans embryos. J Struct Biol (2012). doi:10.1016/j.jsb.2011.12.004

13. Magat, J. et al. 3D MRI of explanted sheep hearts with submillimeter isotropic spatial resolution: comparison between diffusion tensor and structure tensor imaging. Magn Reson Mater Physics, Biol Med (2021). doi:10.1007/s10334-021-00913-4

14. Bernus, O. et al. Comparison of diffusion tensor imaging by cardiovascular magnetic resonance and gadolinium enhanced 3D image intensity approaches to investigation of structural anisotropy in explanted rat hearts. J Cardiovasc Magn Reson (2015). doi:10.1186/s12968-015-0129-x

15. Mulet, A. Book Review: Modern Drying Technology, Volume 3: Product Quality and Formulation, edited by E. Tsotsas and A. S. Mujumdar. Dry Technol (2014). doi:10.1080/07373937.2013.860580

16. Bray, D. Critical Point Drying of Biological Specimens for Scanning Electron Microscopy. in Supercritical Fluid Methods and Protocols (2003). doi:10.1385/1-59259-030-6:235

17. Hall, D. J., Skerrett, E. J. & Thomas, W. D. E. Critical point drying for scanning electron microscopy: a semi-automatic method of preparing biological specimens. J Microsc (1978). doi: 10.1111/j.1365-2818.1978.tb00106.x

18. Bray, D. F., Bagu, J. & Koegler, P. Comparison of hexamethyldisilazane (HMDS), Peldri II, and critical-point drying methods for scanning electron microscopy of biological specimens. Microsc Res Tech (1993). doi:10.1002/jemt.1070260603

19. Nation, J. L. A new method using hexamethyldisilazane for preparation of soft insect tissues for scanning electron microscopy. Biotech Histochem (1983). doi:10.3109/10520298309066811

20. Tran, T. et al. Correcting the shrinkage effects of formalin fixation and tissue processing for renal tumors: Toward standardization of pathological reporting of tumor size. J Cancer (2015). doi:10.7150/jca.12094

21. Jonmarker, S., Valdman, A., Lindberg, A., Hellström, M. & Egevad, L. Tissue shrinkage after fixation with formalin injection of prostatectomy specimens. Virchows Arch (2006). doi:10.1007/s00428-006-0259-5

22. Metscher, B. D. Micro CT for comparative morphology: Simple staining methods allow high-contrast 3D imaging of diverse non-mineralized animal tissues. BMC Physiol (2009). doi:10.1186/1472-6793-9-11

23. Gignac, P. M. & Kley, N. J. Iodine-enhanced micro-CT imaging: Methodological refinements for the study of the soft-tissue anatomy of post-embryonic vertebrates. J Exp Zool Part B Mol Dev Evol (2014). doi:10.1002/jez.b.22561

24. Rajasekar, A., Trew, M. L. & Sands, G. B. Understanding and enhancing the use of micro-computed tomography in soft tissue. (2015). doi:10.17608/k6.auckland.15026040

25. Streeter, J. Gross morphology and fiber geometry of the heart. in Handbook of Physiology: The Heart 61–112 (American Physiological society, 1979).

26. Gilbert, S. H. et al. Visualization and quantification of whole rat heart laminar structure using high-spatial resolution contrast-enhanced MRI. Am J Physiol - Hear Circ Physiol (2012). doi:10.1152/ajpheart.00824.2011

